# High fidelity hypothermic preservation of primary tissues in organ transplant preservative for single cell transcriptome analysis

**DOI:** 10.1101/115733

**Authors:** Wanxin Wang, Lolita Penland, Ozgun Gokce, Derek Croote, Stephen Quake

**Author notes:** Corresponding Author, Stephen Quake, James H Clark Center E300, 318 Campus Drive, Stanford CA 94305. Equal contributors (Wanxin Wang, Lolita Penland).

## Abstract

**Background:** High-fidelity preservation strategies for primary tissues are in great demand in the single cell RNAseq community. A reliable method will greatly expand the scope of feasible collaborations and maximize the utilization of technical expertise. When choosing a method, standardizability is as important a factor to consider as fidelity due to the susceptibility of single-cell RNAseq analysis to technical noises. Existing approaches such as cryopreservation and chemical fixation are less than ideal for failing to satisfy either or both of these standards.

**Results:** Here we propose a new strategy that leverages preservation schemes developed for organ transplantation. We evaluated the strategy by storing intact mouse kidneys in organ transplant preservative solution at hypothermic temperature for up to 4 days (6 hrs, 1, 2, 3, and 4 days), and comparing the quality of preserved and fresh samples using FACS and single cell RNAseq. We demonstrate that the strategy effectively maintained cell viability, transcriptome integrity, cell population heterogeneity, and transcriptome landscape stability for samples after up to 3 days of preservation. The strategy also facilitated the definition of the diverse spectrum of kidney resident immune cells, to our knowledge the first time at single cell resolution.

**Conclusions:** Hypothermic storage of intact primary tissues in organ transplant preservative maintains the quality and stability of the transcriptome of cells for single cell RNAseq analysis. The strategy is readily generalizable to primary specimens from other tissue types for single cell RNAseq analysis.

## Background

Quantitative profiling of transcriptome landscapes at single cell resolution (scRNAseq) has brought new insights in understanding cell types [1,2], states [3], and interactions [4] in the inherently heterogeneous primary tissues. It, however, has also raised new logistical challenges in the specimen conduit from tissue collection sites to laboratories. The low tolerance to cell damage and RNA degradation in scRNAseq and the less resilient nature of cells in primary tissues make it imperative to process primary specimen immediately after procurement, imposing logistical hurdles especially for collaborations in a multi-institutional setting.

A preservation strategy enabling primary tissue storage and transportation will greatly change the status quo and facilitate collaboration between basic science laboratories and distributed medical centers where tissue is collected. Cryopreservation and chemical fixation have been pursued [5-8], but neither is proven ideal for scRNAseq. In the case of cryopreservation in scRNAseq, a recent study reported tolerable impact of elevated cell death from freeze-thaw on cell lines and specimens with well-represented cell types [6]; another study reported an insufficient recovery and reduced transcriptome complexity for low-abundant and less resilient populations [5]. The susceptibility to handling variations and potential variability in freezing media compositions (e.g. serum) pose challenges for standardization. Furthermore, cryopreservation requires mincing the sample as well as maintenance of temperatures down to -80 °C. Crosslinking-based chemical fixation, on the other hand, suffers from low recovery of intact mRNA, while alcohol dehydration-based fixation has yet to show high generalizability from cell cultures to primary tissues [8]. Moreover, for efficient isolation of single cells, fixation is preferentially done on single-cell suspensions [7, 8], making it a necessity to perform tissue dissociation, usually a critical step for scRNAseq, at tissue collection sites. In fact, the aforementioned approaches all require that a multi-step protocol be performed at the collection sites, where experienced personnel are not always available and technical variations can be introduced.

An ideal strategy would avoid drastic physical or chemical changes on the primary specimen and require minimal processing at tissue collection sites. This requirement has much in common with the preservation of organ transplants, which uses hypothermic temperatures to reduce cell metabolism and increase tolerance to insults such as ischemia and hypoxia [9]. Preserving solutions for hypothermic organ preservation hence are designed to address cell-injuring events caused by hypothermia, including ionic imbalance, acidosis, and free radical production [9-11]. Exemplars such as the University of Wisconsin (UW) solution demonstrated high generalizability in preserving post transplant functionality of pancreas (72 hr), kidney (72 hr), and liver (30 hr) [11]. A commercial preparation of such preservative for research use, Hypothermosol-FRS (HTS-FRS), has been increasingly employed in handling primary tissues [12], cells [13-15], and engineered tissue products [16,17]. Comparative studies done on a spectrum of sample types including human hepatocytes [13], coronary artery smooth muscle cells [14], bone-marrow derived mesenchymal stem cells [15], and mouse hippocampus [12] demonstrated superior efficacy of HTS-FRS in maintaining cell viability compared i) to cell culture media and, in some cases, UW solution in hypothermia as well as ii) to cryopreservation. However, previous studies of the physiologic effects of this approach were mainly limited to viability assays and microscopic examination that interrogate membrane permeability, metabolic activity, cell morphology and surface marker expression. Given that the rapid degradation of RNA could precede the deterioration of these examined parameters [18], previous reports are insufficient to conclude whether preservation fidelity is suitable for scRNAseq.

Here we evaluated hypothermic preservation of primary specimen in HTS-FRS for use in scRNAseq. We used FACS to compare viability of cells recovered from fresh and preserved mouse kidneys, and used scRNAseq analysis to demonstrate the efficacy of the strategy in preserving population heterogeneity, transcriptome integrity, and transcriptome stability of kidney resident immune cells in kidneys undergoing up to 3 days of preservation. This approach enables one to preserve intact primary specimen at 4°C for periods suitable for long distance transportation of samples and standardization of experimental approaches in expert labs.

## Results

We designed our experimental procedure such that preservation preceded dissociation (Figure 1). Specifically, intact kidneys were preserved for 0, 6 hour, or 1-4 days immediately following harvest. After the chosen duration of preservation, we enzymatically digested the tissues into single cell suspension and used FACS to assess overall cell viability. We then used the surface marker Cd45 to enrich for kidney resident immune cells to further evaluate our strategy in the context of scRNAseq, given i) that this population encompasses a diverse spectrum of immune lineages with varying abundance, and ii) the population’s critical role in renal injuries and diseases [19-21].

**Figure 1.**
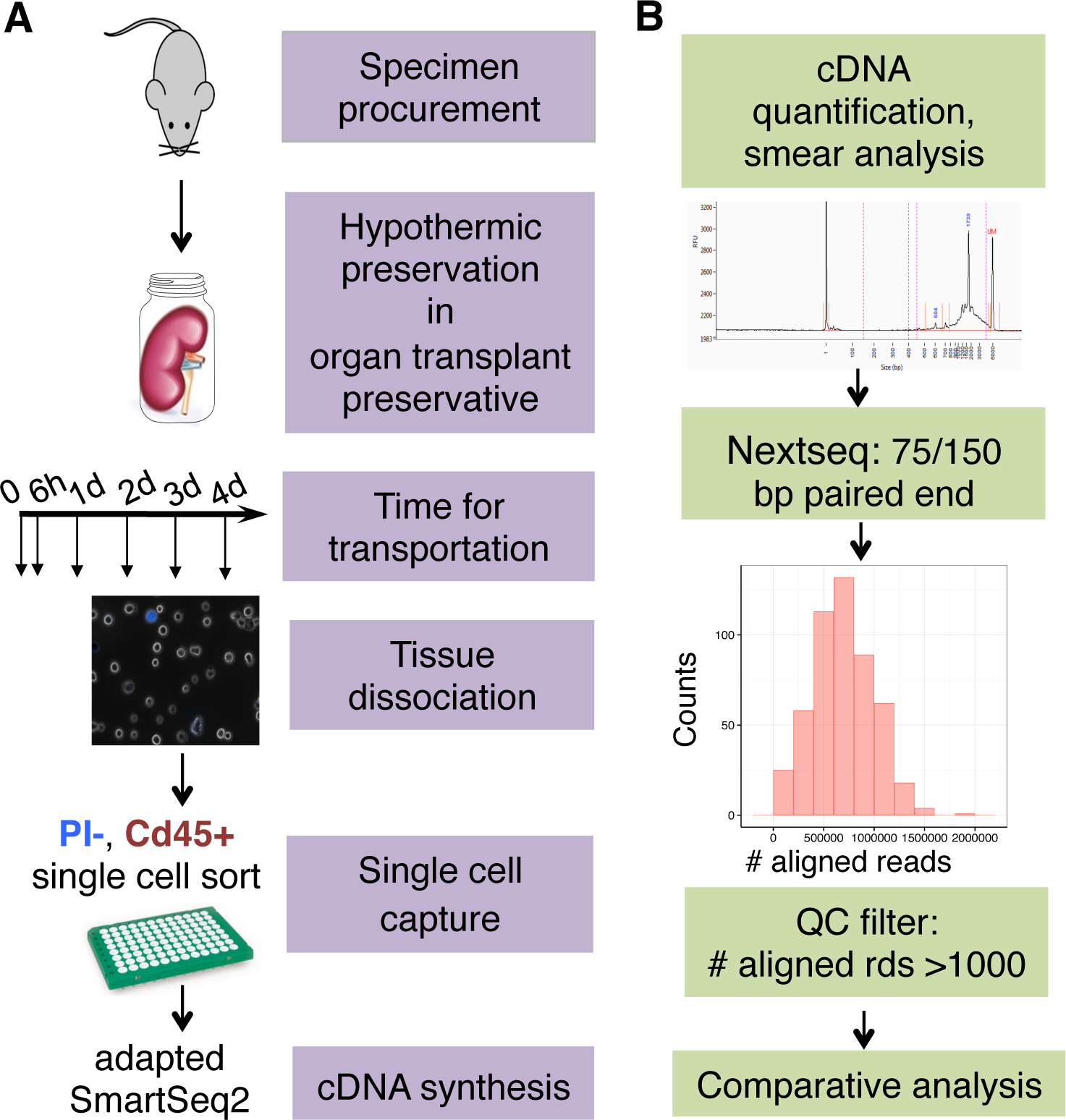
Pipeline design. (A) Schematics and order of preservation, tissue dissociation, single-cell capture, and cDNA synthesis. (B) Criteria for data filtering.

Over the examined durations of preservation, we observed no notable time-associated reduction for the fractions of propidium iodide negative (PI-) and Cd45+ populations (Figure 2A, Additional file 1: Figure S1), suggesting that the strategy effectively retained the overall cell viability and the cell surface marker integrity. For each timepoint, PI- and Cd45+ single cells were sorted for scRNAseq analysis. cDNA sysnthesis on sorted cells gave no notable smearing towards lower fragment sizes for preserved samples (Additional file 1: Figure S2) and comparable success rates in getting sufficient cDNA (≥ 2 ng) between fresh samples and those after up to 3 days of preservation (Figure 2A). The success rate dropped notably at day 4, despite that the fraction of PI- cells stayed comparable with that of fresh, indicative of the notation that mRNA degradation preceded cell membrane permeabilization in early cell death events [18].

**Figure 2.**
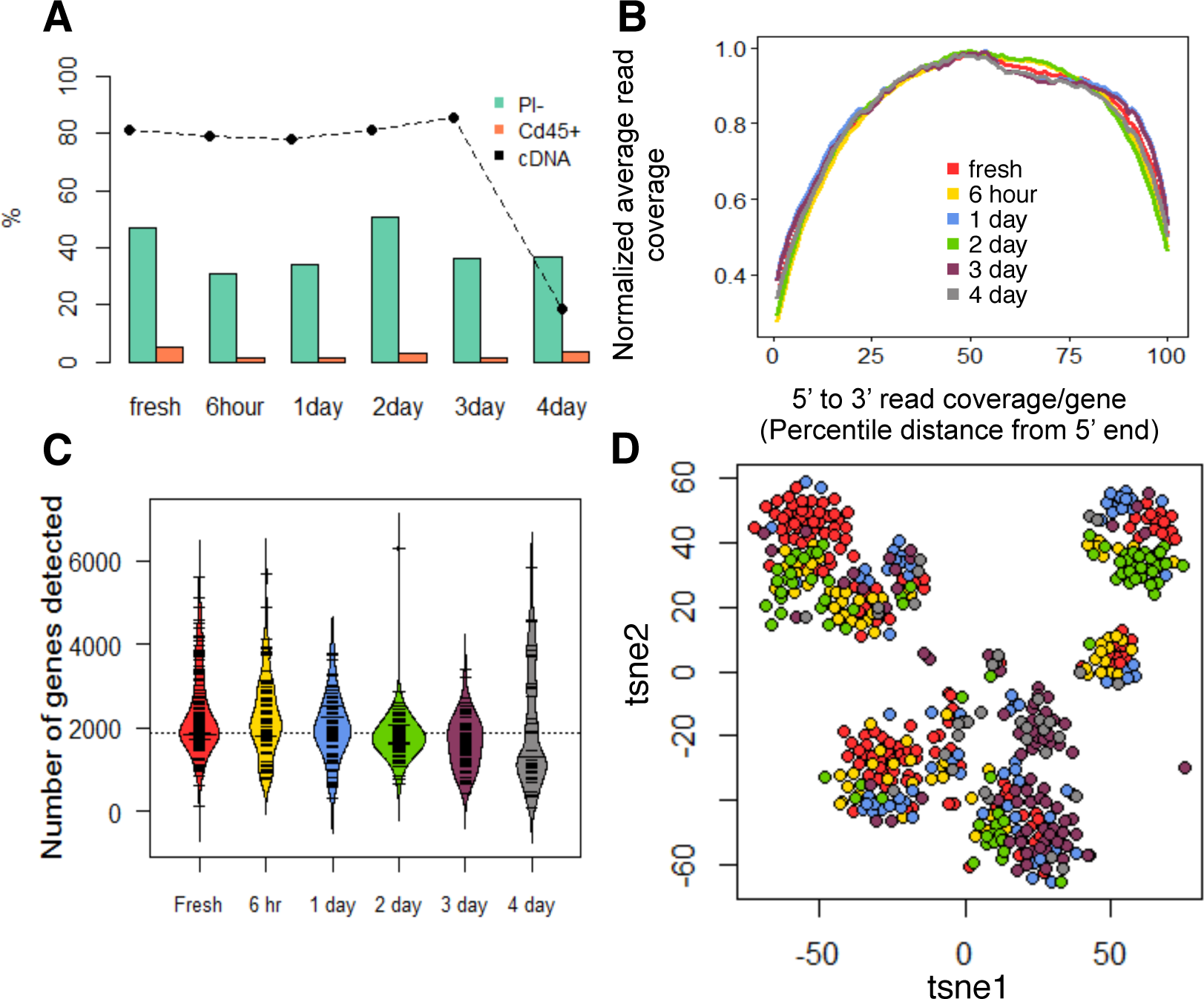
Quality comparison of single cells recovered from fresh and preserved samples (6h-4d) in terms of (A) Overall cell viability (PI-), Cd45+ population abundance, success rates getting sufficient cDNA for sequencing. (B) 5’-3’ read coverage on exons. (C) Distribution of number of detected genes over preservation time. d tSNE on all detected genes. (Coloring in B, C, D all follows legend in B)

510 single cells with sufficient cDNA level (≥2 ng) were sequenced; 502 (98%) passed quality filtering and were retained for downstream analysis (Figure 1B and 2A). To further evaluate mRNA integrity, we examined 5’ to 3’ read coverage across all exons for each single cell, and observed no more bias towards 3’ in preserved samples than in fresh samples (Figure 2B), which was further supported by both qualitative inspection of the coverage curves for individual cells (Additional file 1: Figure S3) and quantitative assessment of the collective skewness of curves for each timepoint (Additional file 1: Figure S4). In addition, the number of genes detected per cell did not drop noticeably until after 4 days of preservation (Figure 2C).

We next assessed the impact of preservation on the cell type heterogeneity of kidney resident immune cells. To explore the data in an unbiased manner, we performed dimensional reduction using whole-transcriptome information via t-distributed stochastic neighbor embedding (tSNE). In the resulting 2-dimensional tSNE space (Figure 2D), single cells formed well-segregated clusters, which we defined into 9 putative clusters computationally (Figure 3A). Given that it is clear that preservation time is not the driving force for the segregation of the clusters (Figure 3B), we hypothesized that the source of the segregation was the cell type heterogeneity in kidney resident immune cells, and performed hierarchical clustering on a panel of canonical markers that define major lineages of immune cells. The resulting outcome of the clustering was nearly identical to that observed in the tSNE space (Figure 3C), confirming our hypothesis.

**Figure 3.**
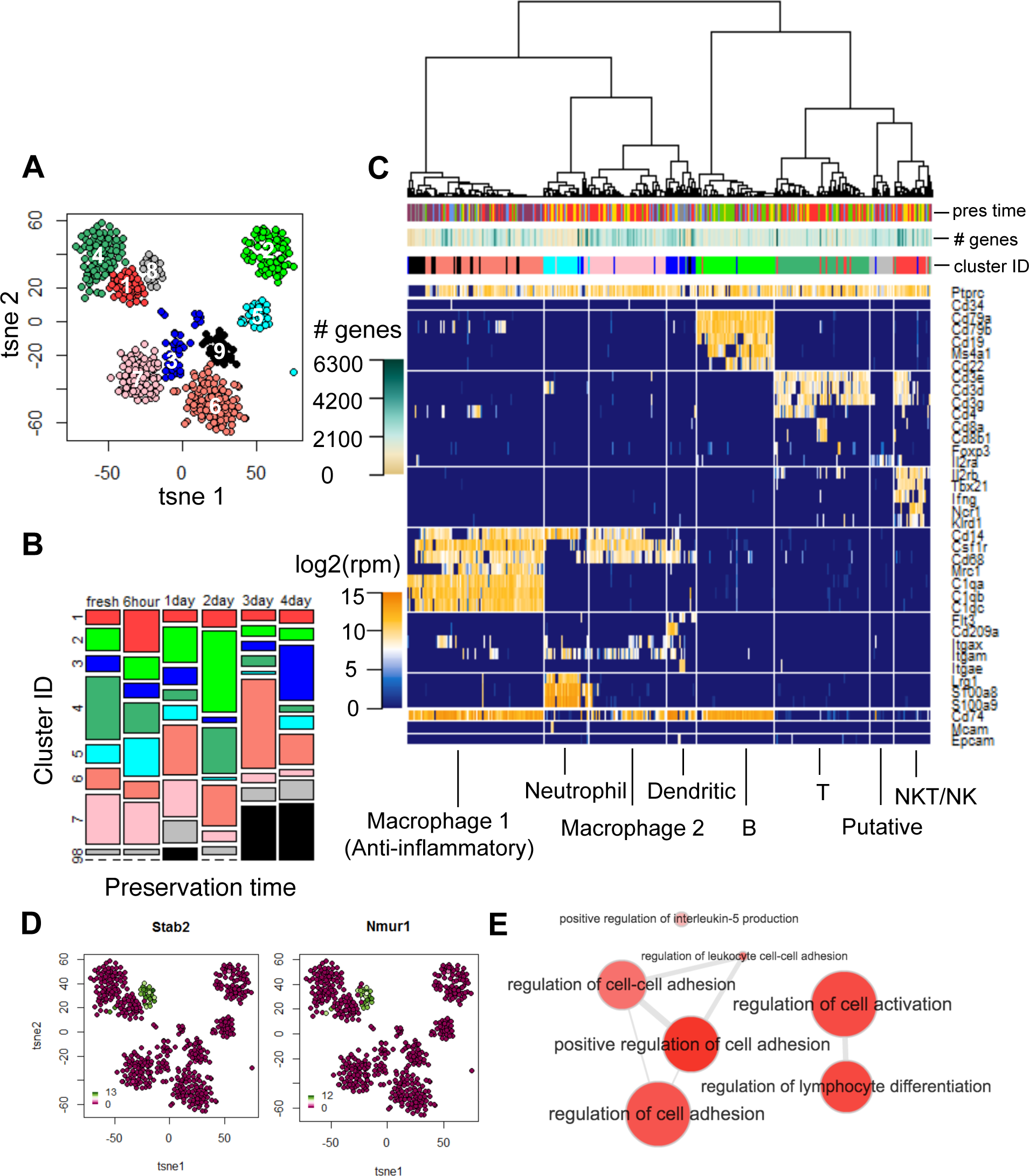
Cell types in kidney resident immune cells and the impact of preservation on cell type heterogeneity. (A) Definition of putative cell clusters on 2d tsne. (B) Distribution of putative clusters in fresh and preserved tissues (6h-4d). (C) Identification of putative clusters with known cell types using hierarchical clustering on a canonical panel of genes defining major immune lineages. (D) Expression of representative genes differentially expressed in cluster 8 (color bar shown in log2(rpm+1)). (E) Hierarchy of enriched functional terms for differentially expressed genes in putative cluster 8. (All coloring and numbering for cluster id in B, C follow those in a; coloring for preservation time follow that in Fig. 2B.)

It is noteworthy that although 7 out of the 9 clusters defined by both methods can be unambiguously identified as known immune populations (Figure 3C), there are 2 (cluster 8 and 9) that cannot be assigned using classical definitions. Unbiased differential expression analysis on cluster 8 revealed a list of genes that are uniquely expressed in this cluster (Figure 3D, Additional file2: Table S1) with enriched ontology terms (Figure 3E, Additional file3: Table S2) suggesting it is a putative lymphocyte population that resembles T cells but lacks classical T cell marker expression. Cluster 9, on the other hand, is most likely a low quality/apoptosing macrophage population due to its absence in fresh samples (Figure 3B), the lack of uniquely expressing markers identified (Additional file1: Figure S5B), the lack of Cd45/Ptprc expression (Figure 3C), and the low number of genes expressed (Figure 3C, Additional file 1: Figure S5A), and hence were excluded in further analysis as a defined population. Each defined population, determined by both methods, is a mixture of cells from fresh tissues and tissues after all examined preservation time (Figure 3B and 3C), revealing no preservation associated batch effect. To summarize, the fact that all defined populations were present in fresh tissues and recapitulated over examined duration of preservation validated our definition of cell types in kidney resident immune cells. Moreover, the proposed preservation strategy effectively maintained the heterogeneity of cell types that exist in varying abundance.

We then compared the transcriptome of fresh and preserved cells within each defined cell type. For all cell types, we calculated pair-wise correlations between cells within and between different preservation conditions. The distribution of correlations was not associated with preservation time (Figure 4A and 4B). Variation in the distributions was dictated by, to greater extent, the heterogeneity within a preservation condition, which also demonstrated no association with time. For lymphocyte populations, we specifically examined the transcripts encoding B cell antibodies (Ig) and T cell receptors (TCR) given their essential roles in B and T cell functions and frequent interrogation by single cell studies. As shown in Figure 4C, expression of key components of Ig and TCR were indistinguishable between fresh and preserved tissues. For B cells, extracting full-length antibody sequence is required for in-depth examination on usage and mutation in variable (V) and joining (J) segments, sequence of complementary-determining region 3 (CDR3), isotypes, as well as affinities. We hence evaluated the success rate in obtaining the information using *de novo* assembly and the impact of preservation on it. From all B cells identified from fresh and preserved tissues, we were able to identify contigs containing complete variable and constant regions for Ig heavy and light chains (Additional file 4: Table S3). The rate of dropout events where only one chain is identifiable is comparably low between fresh and preserved tissues when statistical power holds (Figure 4D). To this end, the strategy conserved the transcriptome profile within identified populations.

**Figure 4.**
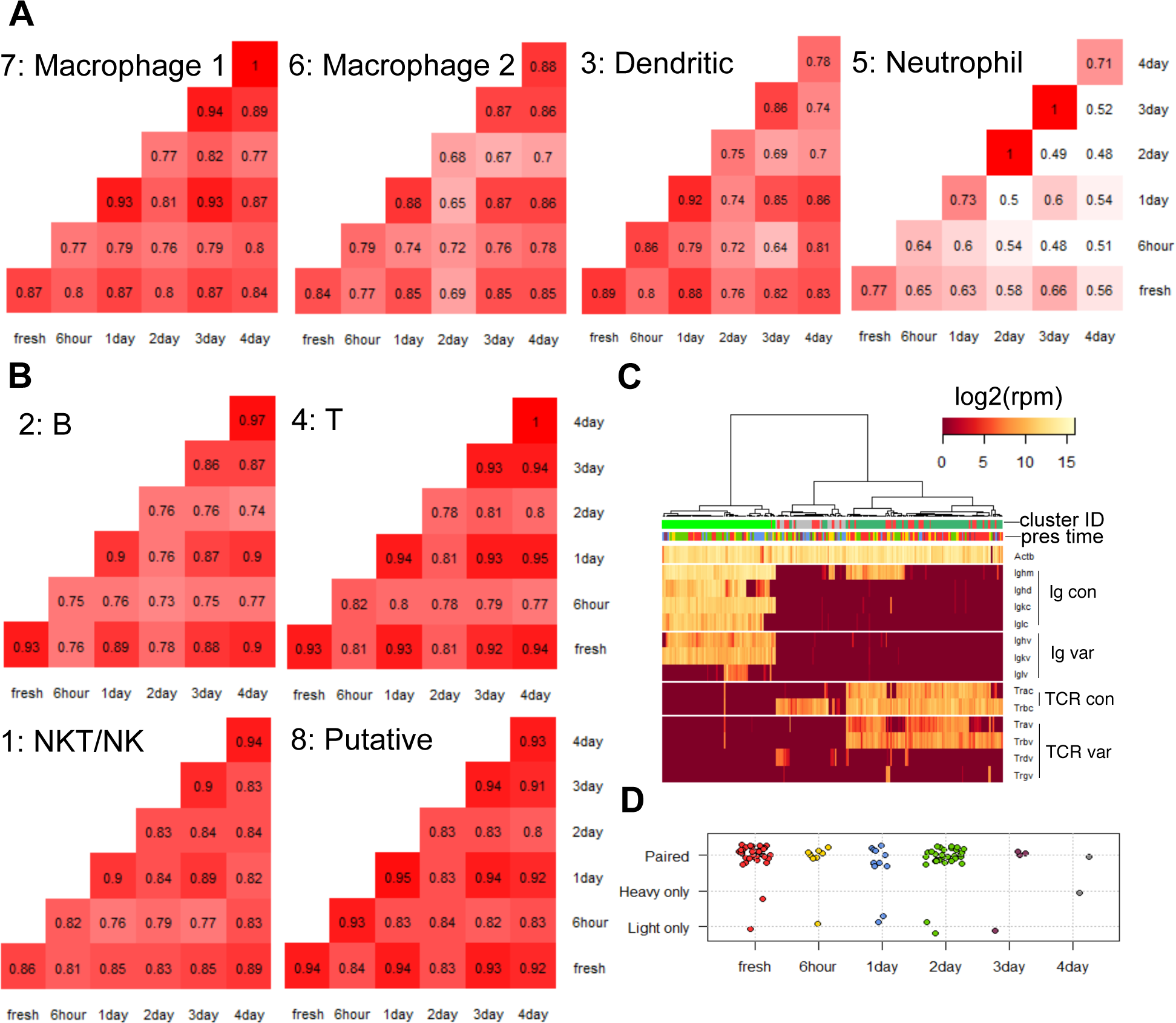
The impact of preservation on the transcriptome profile for each identified cell type. Pair-wise correlation between cells within and across preservation conditions for myeloid (A) and lymphoid (B) populations (Numbers shown are the mediums of each pair of compared distributions. Cell type numbering corresponds to cluster ID in Fig 3A, and cell type identity follows that in Fig 3C). (C) Expression of components of B cell antibodies and T cell receptors in identified lymphoid populations. (D) Distribution of success in extracting full- length transcript sequence for heavy and light chains in all identified B cells. (Coloring for preservation time and cluster ID in C, D follow that in Fig 2B and 3A, respectively.)

## Discussion

Hypothermic preservation of primary tissues in organ transplant preservative effectively maintained the viability, transcriptome integrity, and transcriptome profile stability of cells for scRNAseq. Cells recovered from tissues after up to 3 days of preservation demonstrated minimal 3’ bias in the read coverage of exons and comparable cell type heterogeneity compared to cells from freshly harvested tissues.

Resident immune cells from kidneys were used for evaluation in the context of scRNAseq because of their known heterogeneity and the vast interest they have drawn in kidney injuries. We were able to define 8 cell types in this population, supported by their presence in fresh tissue and consistent recapitulation in preserved tissues. Within each population, the preservation strategy did not introduce quantitative perturbations on the overall transcriptome profile, and faithfully preserved Ig and TCR transcripts to a degree that we could assemble full-length transcript sequences for these highly variable genes for higher resolution interrogations.

This approach enables an actionable time window (48-72 hr) to be opened up for the transportation of primary specimen from the sample collection sites to the technology sites through express couriers.

The strategy is ideal for scRNAseq also due to its high potential in standardization.Preservation can happen on intact tissues immediately after tissue excision with minimum intervention. Procedures that are susceptible to technical variations could be done at the technology sites in a centralized manner, minimizing the introduction of technical noise, especially in highly variable steps such as dissociation. In addition, the preserving solution functions in a serum-free formulation, and hence is free from variations introduced by lot-to-lot difference in serum preparations.

Thanks to the progress made by the organ preservation community, available preservatives such as UW and HTS-FRS already demonstrated high generalizability in preserving functionality in diverse organ types, including pancreas and heart. We therefore expect that the proposed strategy be readily generalizable to other tissue types for scRNAseq as well as for other procedures such xenograft and organoid generation.

## Conclusions

At single cell resolution, primary tissues after 6 hours to 3 days of hypothermic preservation in organ transplant preservative demonstrated similar cell viability, cell type heterogeneity, transcriptome integrity, and transcriptome profile compared to fresh tissues. The strategy is ideal for scRNAseq given its high fidelity and standardizability. The procedure highly resembles the routine handling of specimen in clinics and hence makes it practical to engage clinicians in collaborations, which are essential for the scRNAseq community as well as highly collaborative endeavors such as Human Cell Atlas.

## Methods

### Mouse kidney isolation, preservation, and dissociation

Single-cell experiments were performed on kidneys of CD1 wild type mice. Mice were housed in filtered cages and all experiments were performed in accordance with approved Institutional Animal Care and Use Committee protocols.

Experiments were performed on both lungs or on 1.5 lungs that were pooled from the same mouse for the time points 0-2 days and 3-4 days, respectively. .

Mice of ~3 week old were euthanized by administration of CO_2_. Kidneys were harvested *en bloc* without perfusion and were either dissociated immediately for single cell sort or preserved in the HypoThermosol^®^ FRS solution (BioLifeSolutions) at 4 °C for 6 hrs, 1, 2, 3, or 4 days before dissociation and further processing.

Kidneys were minced with a razor blade and dissociated in Liberase DL (Roche) in RPMI 1640 (LifeTechnologies) with horizontal agitation at 180rpm at 37 °C for 20 min. The resulting single-cell suspension was sequentially passed through a 100 μm, a 70 μm, and a 40 μm strainer (Fisher) and then centrifuged at 300 ×g for 15 min. Pelleted cells were resuspended in ACK red blood cell lysing buffer (Thermo Fisher), incubated for 5 min, quenched with 1 volume PBS (ThermoFisher) containing 2% FBS (ThermoFisher), centrifuged at 300 ×g for 5 min, and then resuspended in FACS staining buffer (BD Biosciences).

### Single-cell sorts, cDNA generation, library preparation, and sequencing

Single cells resuspended in the staining buffer were stained with antibody against surface Cd45 (Cd45-FITC, Sony Biotechnology Inc.) on ice for 20 min following manufacture’s protocol, washed twice with the staining buffer, and then incubated in propidium iodide (PI) solution (Life Technologies) at room temperature for 10 min. Cell viability was evaluated on FACS (Sony Biotechnology Inc.). Singlet PI^-^ Cd45^+^ cells were index sorted onto pre-chilled 96-well plates containing cell lysis buffer using a Sony SH800 sorter.

The plates were vortexed, spun down at 4°C, immediately placed on dry ice, and then stored at -80 °C. Single-cell cDNA libraries were generated using procedures adapted from the SmartSeq2 protocol (Picelli et al. 2014). Briefly, mRNA from single cells in 96- well plates was reverse transcribed using SMARTScribe reverse transcriptase (Clontech), oligo dT, and TSO oligo to generate the first strand cDNA. Resulting cDNA was amplified via PCR (21 cycles) using KAPA HiFi HotStart ReadyMix (KAPA Biosystems) and IS PCR primer. The pre-amplified cDNA was purified using AMPure XP magnetic beads (Beckman Coulter).

Single-cell cDNA size distribution and concentration were analyzed on a capillary electrophoresis-based automated fragment analyzer (Advanced Analytical). Illumina cDNA libraries were prepared for single cells (> 2ng cDNA generated) using Nextera XT DNA Sample Preparation kit (Illumina) with the single cell protocol provided by Fluidigm. Dual-indexed single-cell libraries were pooled and sequenced in 75bp or 150bp pair-ended reads on a Nextseq (Illumina) to a depth of 1-1.5 × 10^6^ reads per cell. CASAVA 1.8.2 was used to separate out the data for each single cell by using unique barcode combinations from the Nextera XT preparation and to generate *.fastq files.

### Processing and analysis of single-cell RNA-seq data

Reads were pre-processed and aligned to mouse reference genome GRCm38 with STAR. For every gene in the reference, aligned reads were converted to counts using HTseq under the setting -m intersection-strict \-s no.

Downstream data analysis was performed in R. Prior to analysis, cells with less than 1000 reads were excluded, reducing the dataset from 510 cells to 502 cells. For each cell, counts were normalized to reads per million (rpm) in log2 scale through division by the total number of aligned reads, multiplication by 1 × 10^6^, and conversion to log with base 2. For tSNE, pair-wise distances between cells were calculated using all genes detected. Dimensional reduction was performed using ViSNE as implemented in the tsne package [22], and subsequent definition of immune cell lineage clusters were done using hierarchical clustering implemented using Ward’s clustering criterion on the resulting two t-SNE dimensions. Differential gene-expression was performed using scde package [23]. Ontology analysis for uniquely expressed genes associated with putative clusters was done using enrichment analysis of biological process for Mus Musculus. Visualization of the hierarchy of the enriched ontology was done using Revigo [24].

### Assembly of B cell antibody heavy and light chains

Full length, paired immunoglobulin heavy and light chain sequences from single B-cells were assembled and annotated by first trimming raw reads with fqtrim, followed by full transcriptome assembly with Bridger [25]. Immunoglobulin contigs identified through the presence of a heavy or light chain constant region sequence were then annotated using IgBLAST [26]. Variable (V), diversity (D), joining (J), and complementarity determining region (CDR) calls were extracted from the IgBLAST output using Change-O [27].

## Abbreviations

scRNAseq: Single cell RNAseq
UW: University of Wisconsin solution **HTS-FRS:** Hypothermosol-FRS
PI-: Propidium iodide negative
tSNE: t-distributed stochastic neighbor embedding
Ig: Immunoglobulin/antibody **TCR:** T cell receptor
V: Variable **D:** Diversity **J**:Joining CDR: Complementarity determining region

## Declarations

### Ethics approval and consent to participate

All animal experiments were performed in accordance with approved Institutional Animal Care and Use Committee (IACUC) protocols authorized by Stanford University.

### Consent for publication

Not applicable.

### Availability of data and materials

The datasets generated and/or analyzed during the current study are submitted to the NCBI Gene expression Omnibus (GEO, http://ncbi.nlm.nih.gov/geo) and will be available upon request.

### Author’s contributions

WW, LP, and SRQ designed the experiments. LP, WW, and OG performed the experiments. WW and DC performed data analysis. WW and SRQ wrote the manuscript.

## Acknowledgements contributions

The author would like to thank Hongxu Ding and Brian Yu for valuable advice and discussion.

## Additional files

**Additional file 1: Figure S1.** Overall viability and abundance of Cd45+ cells across different durations of preservation time. At the bottom of each panel, the percentage in white was calculated as: counts of PI- events / counts of all events for the top panels, and counts of Cd45+ events / counts of all events for the bottom panels. Raw counts were shown in the fractions. **Figure S2.** cDNA concentration and smearing assessed via fragment analysis for single Cd45+ cells from mouse kidneys after different durations of preservation. **Figure S3.** 5’-3’ read coverage on exons for single cells from mouse kidneys after different durations of time. **Figure S4.** Distribution of skewness of 5’-3’ read coverage on exons for single cells from mouse kidneys after different durations of time. **Figure S5.** (A) Number of genes detected cast on 2d tSNE. (B) Uniquely expressing genes identified for each putative cell clusters. (Coloring of cluster ID follows that in Fig 3A.) (PDF 6MB)

**Additional file 2:** Table S1. Differentially expressed genes in the putative cluster (cluster 8). (CSV 5KB)

**Additional file 3:** Table S2. GO ontology of genes differentially expressed in the putative cluster (cluster 8). (CSV 2KB)

**Additional file 4:** Table S3. Detailed annotation on assembled full-length transcripts for antibody heavy and light chains in all identified B cells. (CSV 9KB)

